# Targeted manipulation of abundant and rare taxa in the *Daphnia magna* microbiota with antibiotics impacts host fitness differentially

**DOI:** 10.1101/2020.09.07.286427

**Authors:** Reilly O. Cooper, Janna M. Vavra, Clayton E. Cressler

## Abstract

Host-associated microbes contribute to host fitness, but it is unclear whether these contributions are from rare keystone taxa, numerically abundant taxa, or interactions among community members. Experimental perturbation of the microbiota can highlight functionally important taxa; however, this approach is primarily applied in systems with complex communities where the perturbation affects hundreds of taxa, making it difficult to pinpoint contributions of key community members. Here, we use the ecological model organism *Daphnia magna* to examine the importance of rare and abundant taxa by perturbing its relatively simple microbiota with targeted antibiotics. We used sublethal antibiotic doses to target either rare or abundant members across two temperatures, then measured key host life history metrics and shifts in microbial community composition. We find that removal of abundant taxa had greater impacts on host fitness than did removal of rare taxa and that the abundances of non-target taxa were impacted by antibiotic treatment, suggesting no rare keystone taxa exist in the *Daphnia magna* microbiota but microbe-microbe interactions may play a role in host fitness. We also find that microbial community composition was impacted by antibiotics differently across temperatures, indicating ecological context shapes within-host microbial responses and effects on host fitness.

**Importance:** Understanding the contributions of rare and abundant taxa to host fitness is an outstanding question in host microbial ecology. In this study, we use the model zooplankton *Daphnia magna* and its relatively simple cohort of bacterial taxa to disentangle the roles of distinct taxa on host life history metrics, using a suite of antibiotics to selectively reduce the abundance of functionally important taxa. We also examine how environmental context shapes the importance of these bacterial taxa on host fitness.

## Introduction

The microbes in and on host organism tissue, collectively referred to as the microbiome, are recognized as having important beneficial impacts for the host. Many functions have been tied to bacterial species in the microbiota, including nutrient acquisition for the host^1^ and immune system priming^2^. As most species in host-associated microbiota are difficult to culture, experimental perturbation of the microbiota and subsequent sequencing combined with host fitness metric measurement is a commonly used set of methods to understand functional contributions of microbial taxa to host fitness^3,4,5^. To understand the impacts of individual taxa on host fitness, antibiotics can be chosen to selectively perturb taxa and fitness outcomes can be measured^6^. However, this approach is primarily used in systems with highly complex microbiomes, often with hundreds of interacting taxa impacted by these antibiotics^7,8^. While large-scale perturbations are necessary for understanding overall microbiome structure and broad-level interactions, fundamental questions about host-microbiome interactions can be addressed readily in systems with simpler microbial communities. For example, determining whether host fitness is affected more by overall microbiome diversity (number of distinct taxa) or functional diversity (taxa with distinct functions) is tractable in systems with fewer microbial taxa. Identifying the contribution of numerically abundant taxa to host function is also possible in these systems, as specificity of antibiotic targeting can be greater.

To better understand the relationship between specific taxa in the microbiota and host fitness we applied an antibiotic suppression technique in *Daphnia magna*, a widely used model organism in ecotoxicology^9^, population genomics^10^, and host-parasite dynamics^11^. *Daphnia magna*’s microbiome is relatively simple, with only 10-15 amplicon sequence variants (ASVs) constituting >70% of relative abundance^12,13^. In particular, β-proteobacteria, γ-proteobacteria, and Sphingobacteriia are bacterial Classes consistently identified in the *Daphnia magna* microbiome across environments and genotypes^14,15,16^. Important contributions to host fitness may be directly linked to taxa in these Classes, as removal of the microbiota with broad-spectrum antibiotics has been directly linked to decreases in *Daphnia* growth, survival, and fecundity^17,13,18^. In particular, *Limnohabitans*, a highly abundant genus of β-proteobacteria, has been shown to benefit host fecundity^17^. However, no host fitness benefits have been directly linked to the other bacterial Classes prevalent in the *Daphnia magna* microbiome.

Functions provided by the microbiota to the host can be dependent on biotic and abiotic factors. Environmental factors like temperature^19^, pH^20^, and food availability and diet^21^ alter microbiome composition and gene expression profiles of present taxa. Intrinsic tolerance differences among taxa or host-mediated selection for tolerant taxa may drive changes in community composition, which in turn could influence host fitness. We aimed to investigate this environmental factor-microbiome-host fitness interaction using temperature, because *Daphnia magna* live at a wide range of temperatures^22^ and because temperature influences the *Daphnia magna* microbiota^12,23^. Here, we sought to understand which taxa were affected by environmental change using a cold, environmentally relevant temperature similar to that found in late fall, and whether impacted taxa contributed to host fitness.

To identify taxa in the *Daphnia* microbiota associated with specific host life history traits, we selected antibiotics that each suppressed two of the three major bacterial Classes. We used aztreonam to suppress β-proteobacteria and γ-proteobacteria^24^; erythromycin to suppress γ-proteobacteria and Sphingobacteriia (specifically Bacteroidetes)^25^; and sulfamethoxazole to suppress β-proteobacteria and Sphingobacteriia^26^. We aimed to understand how antibiotic-induced changes to the microbiome impacted host fitness, linking changes in relative abundance to host fitness outcomes. Because the β-proteobacteria *Limnohabitans* impacts host fitness^17^ and is present in high relative abundance in the *Daphnia magna* microbiota, we hypothesized that more abundant taxa contributed a greater share of functions impacting host fitness; as such, suppression of these more abundant taxa would reduce host fitness, specifically in reduced fecundity, survival, and growth. We also aimed to understand how the microbiome and its associated functions changed depending on environmental context. To do this, we raised *Daphnia* in cold temperatures. We hypothesized the differing environment would induce shifts in microbiota composition, as different microbial functions may be necessary to respond to the change. Finally, we combined environmental change with antibiotic treatments to see if reduction of abundant taxa caused differential shifts in host fitness across environments. Here, we hypothesized that reduction of taxa abundant in warmer temperatures with antibiotics would not cause as severe changes in host fitness in the colder temperature treatment, as taxa providing beneficial functions in a different environment would not be targeted by these perturbations and would have reduced competition from the now-suppressed taxa.

## Results

The microbiota of antibiotic-free *Daphnia magna* was dominated by few taxa, with only 10 unique ASVs comprising approximately 60% of total abundance (**Figure 1a**). Primarily, the most abundant ASVs belonged to the *Limnohabitans, Pedobacter*, and *Vitreoscilla* genera. In total, the *Daphnia magna* microbiota was relatively simple, with only 8 bacterial Classes identified (**Figure 1b)**. Of these, Sphingobacteriia and β-proteobacteria were most common (48% and 26%, respectively).

**Figure 1:**
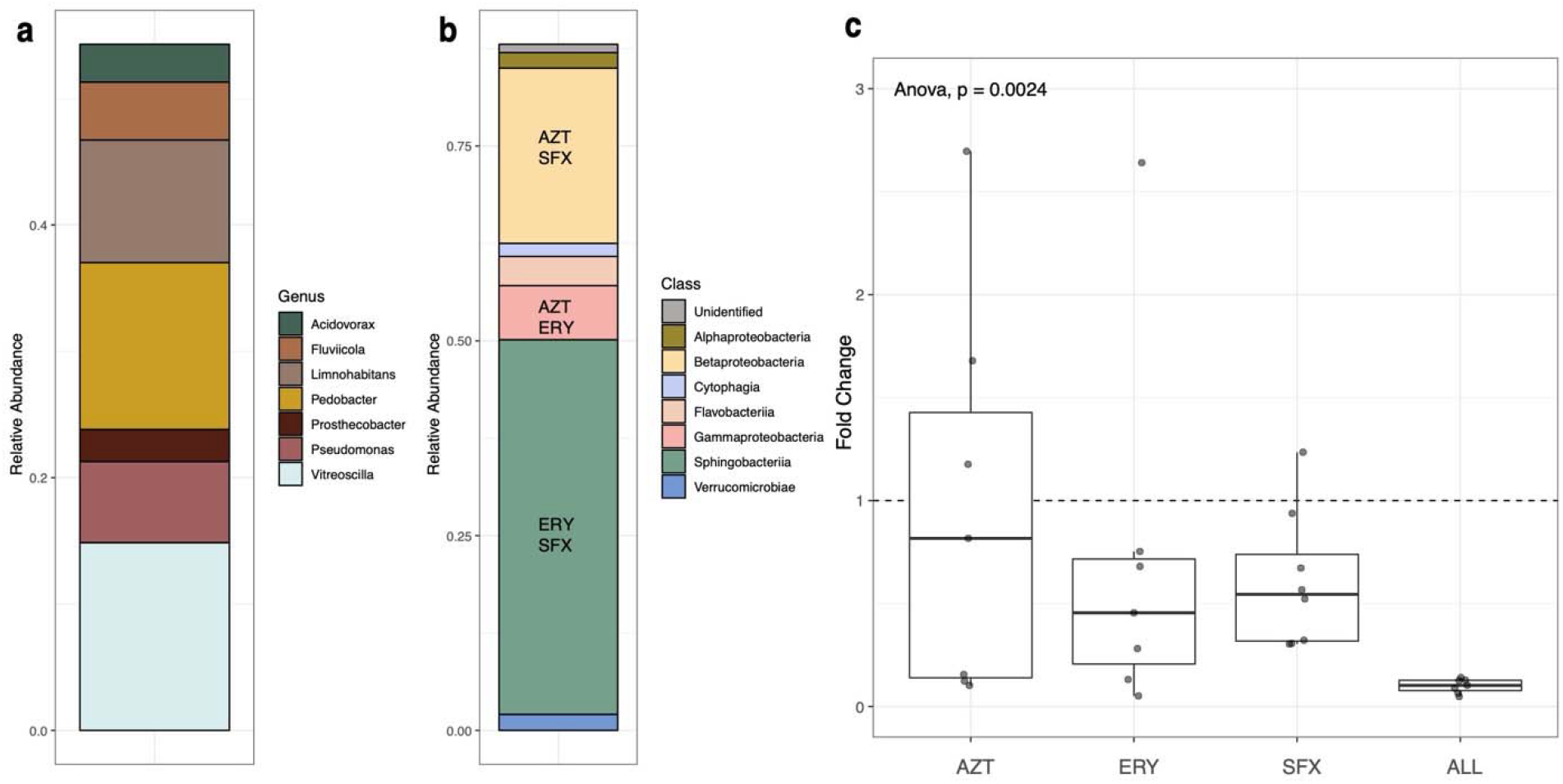
16S rRNA sequencing results from adult *Daphnia magna* in control conditions. (a) Identified genera of the most abundant 10 ASVs in the *Daphnia magna* microbiota and their relative abundances. (b) Relative abundances of bacterial Classes identified in the *Daphnia magna* microbiota. Antibiotics targeting the three bacterial Classes are identified within the columns (AZT = aztreonam, ERY = erythromycin, SFX = sulfamethoxazole). (c) Boxplot of bacterial relative abundance fold change across antibiotic treatments as compared to the control treatment, measured by amplification of the 16S rRNA V4 hypervariable region using qPCR.

Relative abundance of bacteria in *Daphnia magna* was significantly impacted by antibiotic treatment (*F*_4,32_ = 5.197, *p* = 0.0024, **Figure 1c**). Mean fold change in *Daphnia* treated with the antibiotic trio was 0.1-fold that of the control treatment, while treatment with each of the three antibiotics individually slightly reduced relative abundance (AZT 0.96 fold change; ERY 0.71 fold change, SFX 0.61 fold change). The microbiota was significantly impacted by antibiotic treatments at the class rank (PERMANOVA, pseudo-*F*_4,41_ = 3.977, *R*^2^ = 0.26, *p* = 0.01, **Figure 2**). Pairwise PERMANOVA comparisons of antibiotic treatments to the no antibiotic control indicated significant impacts of erythromycin (which suppresses γ-proteobacteria and Sphingobacteriia; *p* = 0.01), sulfamethoxazole (which suppresses β-proteobacteria and Sphingobacteriia; *p* = 0.001), and the antibiotic trio (*p* = 0.001) on microbiota composition (**Supplementary Table 1**). We found multiple ASVs that were differentially expressed in each antibiotic treatment (**Figure 3, Supplementary Table 2**). Though aztreonam-treated samples did not have significantly different overall composition than the control, there were differences in the relative abundance of 8 ASVs, including decreases in *Pseudomonas* (2^−26.5^, or 10^−8^ fewer *Pseudomonas* than in the no antibiotic control) and Sphingomonas (2^−24.3^), and increases in *Microvirga* (2^7.04^). Erythromycin had 16 differentially abundant ASVs, with an unidentified Sphingobacteriia genus and *Emticicia* experiencing the greatest fold abundance changes (2^20.2^ and 2^−9.43^, respectively). Treatment with sulfamethoxazole induced changes for 8 ASVs, primarily increasing the abundance of an unidentified Sphingobacteriia (2^27.4^) and decreasing the abundance of *Pedobacter* (2^−9.19^). Treatment with the antibiotic trio impacted the abundances of 19 ASVs, contributing to fold increases of an unidentified Sphingobacteriia (2^19.24^) and decreasing the abundance of *Caulobacter* (2^−20.18^). In summary, aztreonam, meant to target β-proteobacteria and γ-proteobacteria, reduced the relative abundances of γ-proteobacteria and some α-proteobacteria; erythromycin, meant to target γ-proteobacteria and Sphingobacteriia, increased the relative abundance of α-proteobacteria while decreasing the relative abundances of multiple other Classes including γ-proteobacteria and Sphingobacteriia; sulfamethoxazole, meant to target β-proteobacteria and Sphingobacteriia, decreased the relative abundance of β-proteobacteria and increased α-proteobacteria. The antibiotic trio had the most wide-ranging effects on the microbiota, increasing the relative abundance of some Sphingobacteriia but primarily decreasing relative abundances across multiple Classes.

**Figure 2:**
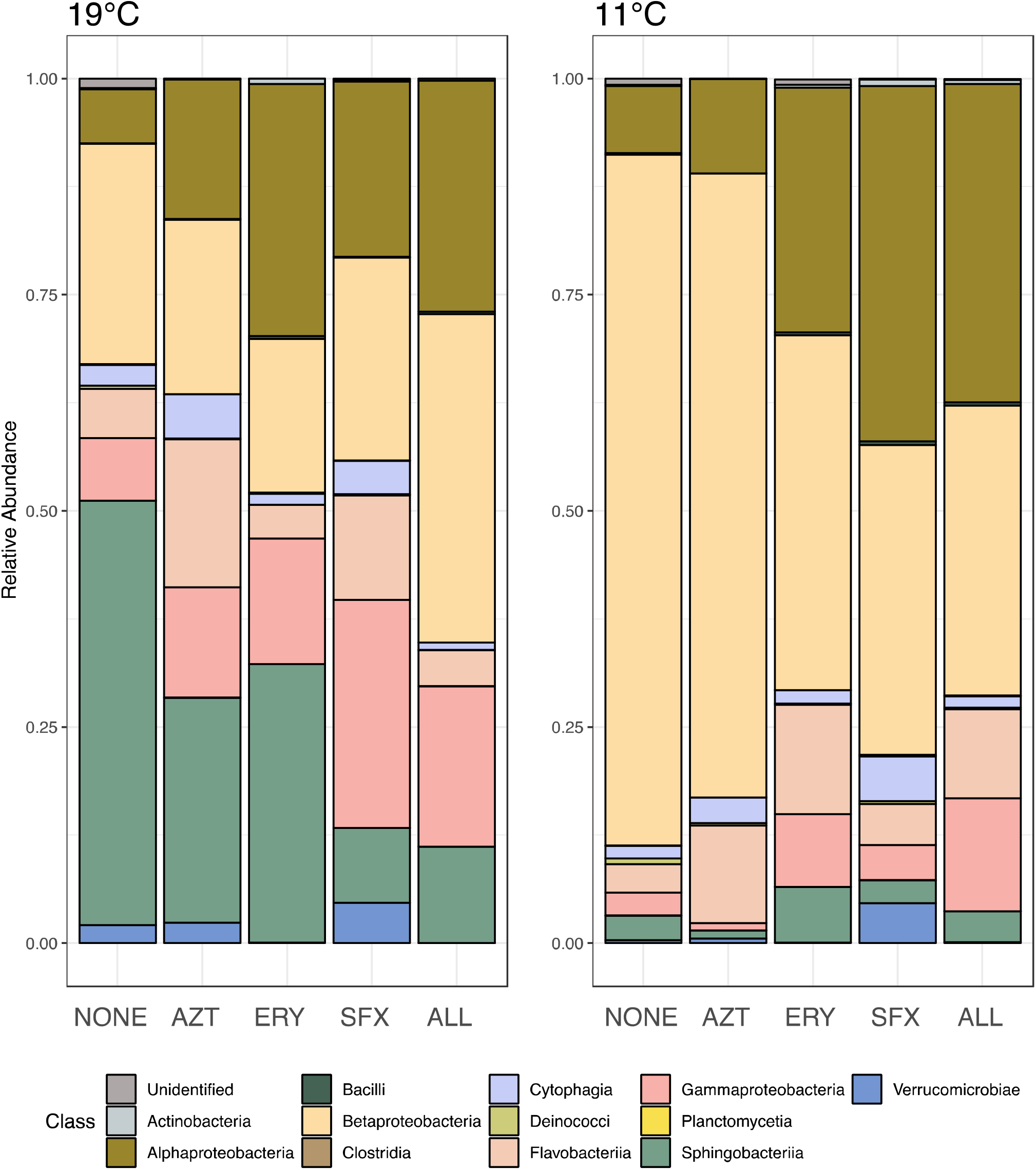
Microbiota composition at the Class level in *Daphnia magna* across antibiotic treatments and across temperature treatments (NONE = no antibiotics, AZT = aztreonam, ERY = erythromycin, SFX = sulfamethoxazole, ALL = AZT, ERY, and SFX). Taxa are conglomerated at the Class rank to show differences in relative abundance of taxa among antibiotic treatments.

**Figure 3:**
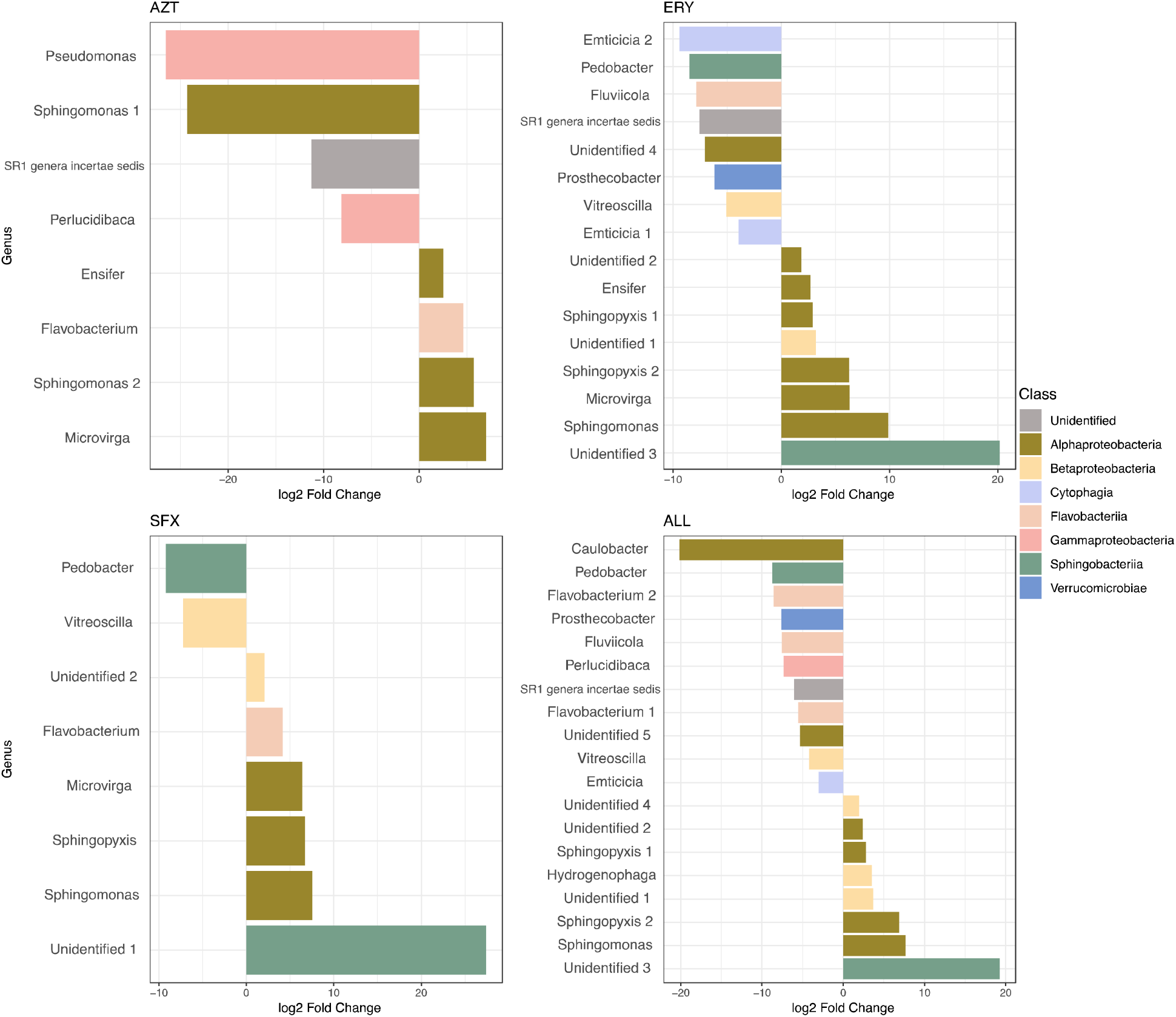
Differentially abundant ASVs in each antibiotic treatment as compared to the no antibiotic control. Each bar represents a single ASV identified to the genus level, with genus name indicated on the left. Bar color indicates the bacterial Class of each ASV, and bar length indicates the fold change in abundance of each ASV.

The *Daphnia magna* microbiota was significantly impacted by temperature. Composition was significantly shifted at the Class taxonomic rank across temperatures (pseudo-*F*_1,41_ = 7.178, *R*^2^ = 0.12, *p* = 0.01) and across the interaction between antibiotic treatment and temperature (pseudo-*F*_4,41_ = 1.623, *R*^2^ = 0.11, *p* = 0.02, **Figure 2**). In all antibiotic treatments except aztreonam, there were significant differences in microbiota composition between temperatures when compared to the control (Pairwise PERMANOVAs, erythromycin pseudo-*F*_1,9_ = 4.001, *R*^2^ = 0.31, *p* = 0.006; sulfamethoxazole pseudo-*F*_1,9_ = 2.605, *R*^2^ = 0.30, *p* = 0.045; trio pseudo-*F*_1,6_ = 5.142, *R*^2^ = 0.46, *p* = 0.034). Multiple taxa were found to be differentially abundant even in treatments not found to have significantly different community compositions overall (**Supplementary Figure 1, Supplementary Table 2**). The no antibiotic control had a single ASV (Sphingobacteriia genus, 2^−13.68^) that was differentially abundant in cold temperatures. In aztreonam, 6 ASVs were reduced in relative abundance in cold temperatures, with a Sphingobacteriia (unidentifiable beyond Class) and an α-proteobacteria genus experiencing the greatest reductions (2^−11.98^ and 2^−11.4^, respectively). In erythromycin, 12 ASVs were differentially abundant; of those, the same Sphingobacteriia as in the aztreonam treatment was reduced (2^−27.54^) and a *Pseudomonas* ASV was more abundant (2^6.34^). Sulfamethoxazole impacted 13 ASVs, including *Limnohabitans* (2^− 6.61^) and the same Sphingobacteriia (2^−25.84^). The antibiotic trio treatment only had one differentially abundant ASV (*Nubsella*, 2^7.89^). Generally, differentially abundant Classes were limited to the α- and β-proteobacteria, Flavobacteriia, and Sphingobacteriia. Sphingobacteriia exhibited the greatest changes in abundance across antibiotic treatments, with a Sphingobacteriia genus experiencing between 2^−5^ and 2^−27.5^ reduced abundance in no antibiotics, aztreonam, erythromycin, and sulfamethoxazole at low temperatures.

Host fitness was significantly impacted by antibiotics. In particular, cumulative host reproduction over the course of the experiment was reduced by antibiotics (*F*_4,470_ = 49.59, *p* < 0.001). Post-hoc Tukey tests revealed that this reduction was most significant in the aztreonam, sulfamethoxazole, and the antibiotic trio treatments (all *p* < 0.001) (**Figure 4a**). A complete list of post-hoc comparisons for cumulative reproduction can be found in **Supplementary Table 3**. Though cumulative host reproduction was reduced in these treatments and many amplicon sequence variants experienced shifts in relative abundance across treatments, there was no single bacterial genus that shifted in relative abundance predictably across treatments, suggesting that there is no genus that is uniquely important to *D. magna* reproductive fitness. Reproductive timing was also impacted (*F*_4,222_ = 4.797, *p* < 0.001), where *Daphnia magna* treated with sulfamethoxazole experienced a later age at first reproduction than those treated with other antibiotics (Tukey HSD, *p* = 0.03, **Figure 4d, Supplementary Table 4**). *Daphnia magna* exposed to antibiotics experienced a significant overall reduction in growth (ANOVA, *F*_4,419_ = 2.08, *p* = 0.004, **Supplementary Table 5**); the main contributor to this was a significant reduction in growth in sulfamethoxazole as compared to erythromycin (Tukey HSD, *p* = 0.003, **Supplementary Table 5**). Exposure to any antibiotic had no impact on *Daphnia* survival (**Figure 4b, Supplementary Figure 2**). Host fitness was also impacted by temperature. Cumulative reproduction was reduced almost completely in the cooler 11°C treatment; no reproduction was observed in the control and only one adult individual reproduced across all of the antibiotic treatments (ANOVA, *F*_1,470_ = 1751.08, *p* < 0.001). There was also an effect of the interaction between antibiotics and temperature (ANOVA, *F*_4,470_ = 19.84, *p* < 0.001), to the extent that the impacts of antibiotics on reproduction in low temperatures was not noticeable due to the strong effect of temperature on reproduction. *Daphnia* growth was limited in the cold temperature treatment (*F*_1,419_ = 176.01, *p* < 0.001) (**Figure 4c**). Temperature impacted survival, with *Daphnia* in colder temperatures surviving significantly more than those in control temperatures (HR = −1.087, *p* = 0.013) (**Figure 4b, Supplementary Figure 2**).

**Figure 4:**
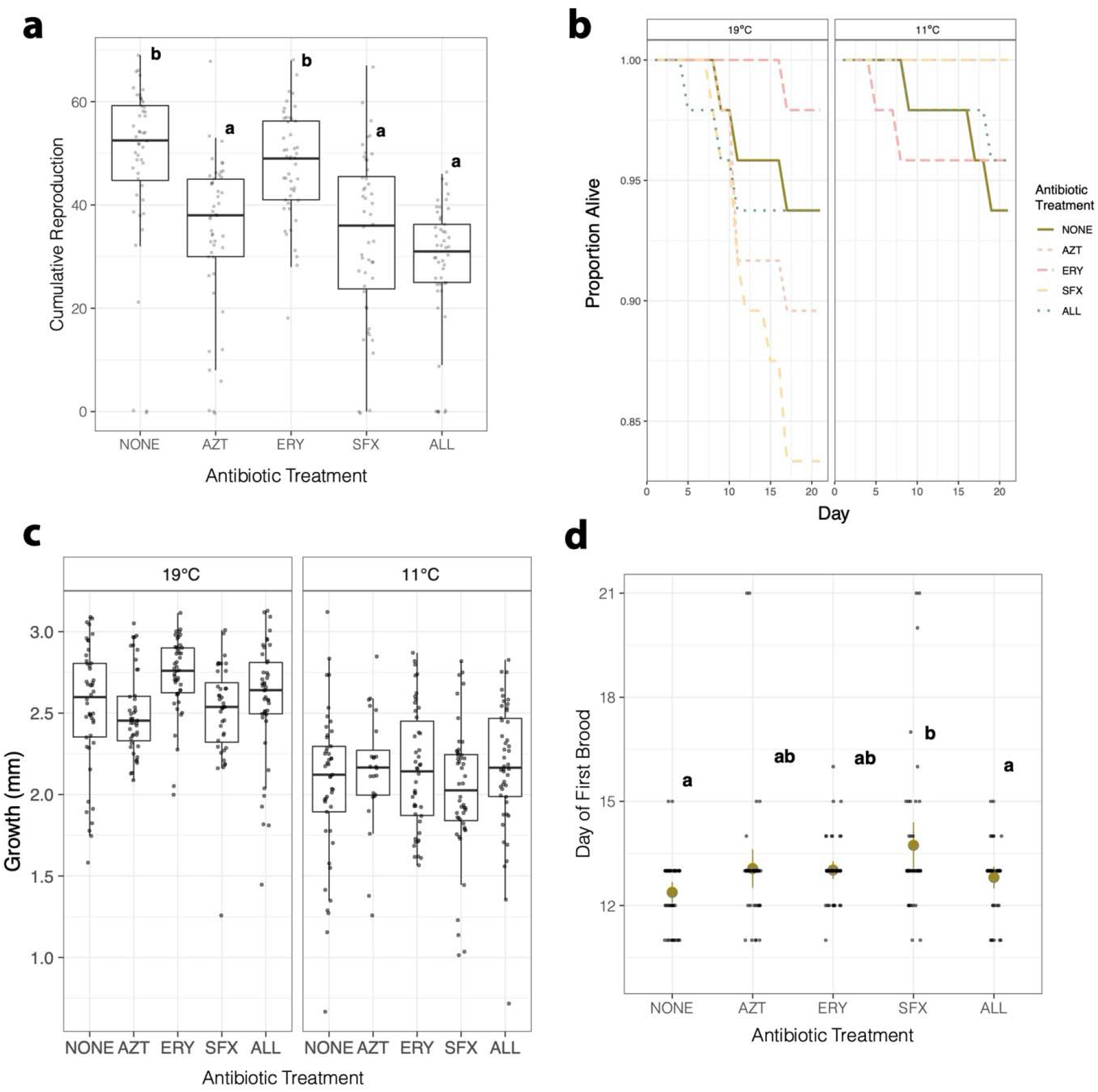
Summary of *Daphnia magna* fitness. (a) Boxplot of cumulative reproduction over the 21-day experiment in *Daphnia magna* across antibiotic treatments (NONE = no antibiotics, AZT = aztreonam, ERY = erythromycin, SFX = sulfamethoxazole, ALL = all three antibiotics) in the 19°C temperature treatment. Points show cumulative reproduction of each individual *Daphnia* over the 21-day experiment. Letters denote significant differences among treatments. (b) Survival curves of *Daphnia magna* in antibiotic treatments across temperature treatments. Line color and pattern denote antibiotic treatment. (c) *Daphnia magna* growth in millimeters over the 21-day experiment across antibiotic and temperature treatments for individuals who survived the entire time course. Boxplots denote median and first and third quartiles, and jittered points show growth of each individual within a treatment. (d) Time to reproductive maturity of *Daphnia magna* in the 19°C temperature treatment across antibiotic treatments. Jittered points denote individuals within each antibiotic treatment. Letters denote significant differences among treatments.

## Discussion

In this study, we manipulated the microbiome of *Daphnia magna* using low doses of targeted antibiotics to examine the impacts of selective suppression on host fitness. We found that aztreonam and sulfamethoxazole, antibiotics targeting abundant bacterial Classes in the *Daphnia magna* microbiome, had the largest impacts on host fitness in both fecundity and growth. However, our results contrast with those of Sison-Mangus *et al*. (2015)^18^ and Callens *et al*. (2016)^14^ in that survival was not impacted by antibiotics here. This is likely due to our use of targeted antibiotics, rather than the broad-spectrum antibiotics used to completely suppress the microbiota in those studies. We found that in all of the antibiotic treatments, relative abundance of bacteria was lower than in the control treatment. However, relative abundance in treatments did not fully correspond with the relative abundances of the intended target groups, suggesting that non-target groups may have also been affected. Alternatively, the *Daphnia magna* microbiota may have a carrying capacity, where non-target groups may replace taxa negatively affected by the antibiotic perturbation. Our 16S rRNA sequencing shows that non-target Classes were impacted in unexpected ways, potentially due to microbial interactions or off-target antibiotic effects. We also found that shifts in microbiome composition were dependent on environmental conditions, with *Daphnia magna* exposed to the same antibiotics at low temperatures not experiencing the same shifts in microbiome composition and differentially affecting host fitness.

Our characterization of the *Daphnia magna* microbiome in standard conditions (19°C, no antibiotics) yields a composition that is similar to that of healthy adult *Daphnia magna* in other studies, though this genotype has been isolated in laboratory culture for >3 years^15,34,37^. This suggests that our *Daphnia magna* maintained in culture have retained *Daphnia*-specific microbes present during initial field collection, as all culture media is autoclaved prior to use and algae is grown axenically. We did find that the microbiota of untreated adult *Daphnia magna* in these cultures exhibited a higher relative abundance of Sphingobacteriia than in other studies (approximately 48%, compared to 4%-20%), but other abundant bacterial Classes were similar, including α-proteobacteria, β-proteobacteria, and γ-proteobacteria^15,34^. In particular, we found that species in the β-proteobacteria genus *Limnohabitans* are highly abundant in the *Daphnia* microbiome, consistent with prior work^15,27,28,29,30^. *Limnohabitans* has important effects on host fitness in *Daphnia magna*, with monoassociations with *Limnohabitans* strains increasing *Daphnia* fecundity^17^. Though species in the *Limnohabitans* genus are prevalent free-living freshwater bacterioplankton^31^, metagenome-assembled *Limnohabitans* genomes from the *Daphnia magna* microbiota have <90% average nucleotide identity shared with their closest sequenced free-living relatives^32^, indicating these *Limnohabitans* species are likely specific to *Daphnia magna*. These *Daphnia*-associated *Limnohabitans* species encode for biosynthesis and export of amino acids essential for *Daphnia magna*^33,34^, potentially providing these when hosts are unable to acquire them from their diet. Other genera identified here and in other studies include *Pedobacter, Emticicia*, and *Acidovorax*^35,15^, among others.

Antibiotics suppressed the microbiota of *Daphnia magna* but were not as targeted as expected. Treatments of aztreonam and sulfamethoxazole reduced the relative abundance of β-proteobacteria as expected; however, erythromycin treatment unexpectedly reduced β-proteobacteria relative abundance as well. Treatment with erythromycin also increased the relative abundance of some Sphingobacteriia ASVs, though it reduced the abundance of Bacteroidetes as expected. This suggests two possibilities: narrow-spectrum antibiotics may be affecting different taxa than expected here, or suppression of the targeted taxa impacts non-targeted taxa that may be interacting with those that were suppressed.

Host fecundity was impacted by treatment with aztreonam, sulfamethoxazole, and the antibiotic trio, and host growth was reduced when treated with the trio as well. Sulfamethoxazole (targeting β-proteobacteria and Sphingobacteriia) also delayed age at first reproduction, supporting our hypothesis that suppression of more abundant bacterial Classes would have larger impacts on host fitness. While it is possible that the antibiotics used here have direct effects on host fitness^30^, germ-free *Daphnia magna* treated with the same antibiotic trio had the same fitness metrics as the *Daphnia* treated here (Cooper, unpublished data), suggesting that differences in host fitness are mediated through the microbiome. The reduction in host fecundity without a consistent associated decrease in abundance of any particular microbial taxon across treatments suggests that multiple taxa are involved in the functions that benefit host fecundity, supported by potential cross-species interactions identified through shotgun sequencing^32^. Specifically, metagenome sequencing indicates that a *Pedobacter* species uniquely encodes for chitin degradation and sialic acid cleavage and that other species (primarily *Limnohabitans*) may be able to utilize those cleaved sialic acids for amino acid biosynthesis^33^. This delay in reproductive maturity and associated reduction in *Pedobacter* in the sulfamethoxazole treatment may indicate that microbe-microbe interactions are affected by targeted antibiotic treatment.

Interestingly, γ-proteobacteria are found in high relative abundances in the *Daphnia magna* gut and could play a role in nutrient acquisition or pathogen protection^14,15^, yet their suppression with erythromycin did not affect host fitness. We hypothesize that this may be due to functional redundancy of taxa found in the *Daphnia* gut, as *Daphnia magna*’s indiscriminate filter feeding exposes gut microbes to a wide array of nutrients^14,36^. Indeed, the abundances and identities of taxa in the *Daphnia* gut vary substantially across studies^23,18,16,30^; this variation may allow different taxa not targeted by erythromycin to retain the necessary functions for nutrient acquisition.

Temperature dramatically shifted the microbiome and the fitness of *Daphnia magna*. These changes have been documented across host genotypes and warmer temperatures^23,12^, but to our knowledge this is the first to examine the effects of this cold of a temperature (11°C) on the microbiota of this keystone species. *Daphnia magna* raised in cold temperatures survived more, grew less, and had almost no offspring, a well-studied physiological mechanism for actively surviving winter in aquatic ecosystems^37^. Correspondingly, the microbiome shifts during this time. In 11°C, β-proteobacteria became even more abundant, comprising >80% of relative abundance. Sphingobacteriia, Flavobacteriia, and γ-proteobacteria were reduced to <5% relative abundance each. This may be due to cold-induced changes to host metabolic processes like fat storage and processing, which have been shown to shape microbiota composition^38^. *Daphnia magna* reduce stearic acid formation at low temperatures but increase monounsaturated fatty acid formation^39^, which could select for taxa able to utilize these types of fatty acids. Alternatively, β-proteobacteria may be so important for host fecundity^17^ that they must remain in high abundance to ensure they remain for the post-winter reproductive cycle. Though microbiota composition shifted in cold temperatures, it is unlikely that host fitness is mediated by microbiota change. Even with a greater relative abundance of reproductive fitness-promoting β-proteobacteria, *Daphnia magna* in cold temperatures had significantly reduced reproductive fitness, suggesting that temperature directly impacts fitness.

Antibiotics did not affect microbiome composition in cold temperatures as they did in standard conditions. Treatment with aztreonam at 11°C resulted in a microbiome composition nearly identical to that of *Daphnia magna* not treated with antibiotics in 11°C. Though erythromycin does not target β-proteobacteria, this Class was reduced in the cold temperature-erythromycin treatment, suggesting that taxa in the *Daphnia* microbiota may be differentially susceptible to antibiotics depending on environmental factors and based on host physiological responses to the environment. Horizontal gene transfer could play a role in this differential response, as species in the *Daphnia magna* microbiome do encode for antibiotic resistance and efflux^33^ and this experiment was conducted over a time period long enough to allow antibiotic resistance to establish within species^40^. Furthermore, cold temperatures have been shown to increase the abundance of antibiotic resistance genes^41^.

Our results suggest that more abundant Classes in the microbiome (in this case, Sphingobacteriia and β-proteobacteria) have larger impacts on host fitness than rarer taxa. Within-host microbial communities generally have a skewed abundance pattern, where a few species constitute the majority of total abundance but many species are found in low abundances. Some work indicates that abundant taxa contribute to host fitness^42^, while others indicate that rare, keystone taxa have disproportionate impacts on host fitness^43^. However, a general relationship between abundance in the microbiota and benefit to host fitness is hard to untangle in complex systems, and much research in model systems focuses on hosts with single microbial taxa that have significant impacts on host fitness^44^. Utilizing animal models like *Daphnia magna* with more than one taxon contributing to host fitness but a relatively simple overall microbial community allows for a greater understanding of the interplay between the microbiota and host. In *Daphnia magna*, more abundant taxa (e.g., *Limnohabitans*) confer greater benefits to host fitness, primarily through functions that contribute to increases in host fecundity and growth, whereas the loss of rare species had little effect on fitness. At the same time, abiotic conditions can have a much larger effect on host fitness than the microbiome, as demonstrated by the changes in *Daphnia* fitness with temperature not directly mirrored by changes in the microbiota. Our results show that multiple members of the *Daphnia magna* microbiota impact host fitness in different ways, and that these impacts must be understood in the broader context of external factors known to directly affect host fitness.

## Methods

### Daphnia

This experiment was conducted using *Daphnia magna* clone 8A, taken from Kaimes Farm, Leitholm, Scottish Borders^45^. Stock cultures of *D. magna* clone 8A were maintained in 19°C controlled chambers with a 16h light, 8h dark light cycle in 400 mL jars with phosphorous- and nitrogen-depleted COMBO medium^46^ for multiple generations. Cultures were fed a standardized 0.25 mg C/mL/day using green algae *Chlamydomonas reinhardtii* (CPCC 243). *C. reinhardtii* was cultured in COMBO medium. The volume necessary to provide *D. magna* with adequate carbon was calculated using the Biotek Epoch Microplate Spectrophotometer.

### Experimental design

Prior to the experiment, 72 *D. magna* were moved to 35 mL glass vials with COMBO medium and allowed to mature under controlled conditions. Neonates from the third brood of each adult were pooled within 24h of birth and randomly assigned to experimental treatments (n=48 per experimental treatment). The experimental treatments consisted of five antibiotic treatments crossed with two temperature treatments, 19°C and 11°C. Antibiotic treatments were as follows: a control treatment with no antibiotics; 500 ug/L aztreonam; 400 ug/L erythromycin; 250 ug/L sulfamethoxazole; and an antibiotic trio consisting of all three antibiotics together at the concentrations listed above. These antibiotic concentrations were chosen on the basis of a pilot experiment showing no short-term toxicity effects on *D. magna* survival but a significant reduction in bacterial abundance (revealed using qPCR with universal bacterial 16S primers). Body size of each *D. magna* was measured from eyespot to beginning of apical spine before placement into the experimental vials. Experimental *D. magna* were raised in 35 mL glass vials with COMBO for 21 days. Each vial was checked for survival and fed daily with 0.25 mg C/ml/day of the diet treatment. Vials were also checked daily for offspring, which were counted and removed if present. At the conclusion of day 21 or upon death, *D. magna* were collected from the treatments. Body size was again measured from eyespot to beginning of apical spine to determine growth. *D. magna* from each treatment were pooled in sets of 10 in 1.5 mL microcentrifuge tubes for DNA extraction and processing (n=4 per treatment).

### DNA extraction, library preparation, and sequencing

DNA was extracted from all pooled samples using the Qiagen DNEasy Blood & Tissue Kit using the manufacturer’s spin-column protocol of total DNA from animal tissues (Qiagen, Hilden, Germany). Whole *D. magna* were digested with Proteinase K for 24h to ensure cells within the carapace were lysed but the carapace was not^47^. Following extraction, PCR amplification of the V4 region of the 16S rRNA gene was performed using the 515f (5’-GTGCCAGCMGCCGCGGTAA-3’) and 806r (5’-GGACTACHVGGGTWTCTAAT-3’) universal 16S primer pair^48^. Amplification consisted of denaturation at 95°C for 3min, followed by 35 cycles of 95°C for 45sec, 58°C for 30sec, and 72°C for 45sec, and finished with a 72°C for 5min extension step. Simultaneously, a subset of samples from each of the antibiotic treatments was prepared for qPCR using the FastStart SYBR Green Master Mix to verify antibiotic treatments were reducing overall bacterial abundance. Each sample was run in triplicate to ensure amplification was achieved in each sample. All samples were checked for successful amplification using a 1% agarose electrophoresis gel. Samples were then normalized with the SequalPrep Normalization Plate Kit. Prior to sample pooling, sample quality was checked using the Agilent High Sensitivity DNA Kit on the Agilent TapeStation and via qPCR with the KAPA Library Quantification Kit. Samples were then pooled and spiked with PhiX DNA. The pooled libraries were then sequenced using the Illumina MiSeq Reagent Kit v2 (300-cycles) on an Illumina MiSeq. Sequencing was carried out at the Nebraska Food for Health Center (Lincoln, Nebraska, USA).

### Sequencing data processing

Following sequencing, reads were demultiplexed using Illumina’s built-in MiSeq Reporter software. All reads were then analyzed using DADA2^49^ in R. In DADA2, our pipeline consisted of low-quality (<Q30) read trimming, estimation of read error, dereplication of reads within samples, and chimera removal. Remaining reads were considered amplicon sequence variants (ASVs), then were assigned taxonomy to the genus level using the Refseq-RDP database^50^. All visualization of ASVs was performed with Phyloseq^51^ in R, where reads without a taxonomic assignment at the phylum level and those assigned to “Chloroplast” were removed for visual clarity. All scripts for read processing and visualization are available on GitHub (https://github.com/reillyowencooper/ab-targeting-daphnia).

### Statistical analysis

All statistical tests were performed in R. Host *D. magna* life history traits measured as indicators of fitness outcomes included growth, survival, and reproduction. Growth was quantified as the difference between size measurements at the beginning and end of the experiment. Differences in growth among treatments, including the interactions between antibiotics and temperature, were analyzed using an ANOVA. We also used ANOVAs to test for effects of antibiotics and temperature on reproduction, which was measured as number of juveniles per brood and day of first reproductive event (production of the first brood for each individual). Tukey’s HSD post-hoc tests were conducted to determine which treatments significantly differed from the control treatment. Survival rates among treatments were analyzed using the Cox proportional hazards model. Individuals alive at experiment conclusion were coded as censored. We used the ΔΔCt method to calculate log fold change in abundance of the 16S rRNA gene among antibiotic treatments, normalizing against the *Daphnia magna* actin gene (forward primer: 5’-CCACACTGTCCCCATTTATGAA-3’, reverse primer: 5’-CGCGACCAGCCAAATCC-3’) and against the control treatment. A PERMANOVA was conducted among treatments on the calculated unweighted UniFrac distances to test the effects of antibiotics and temperature on microbiota composition, then pairwise comparisons of antibiotic treatments in each temperature were conducted to find treatments with significantly different overall community composition. DESeq2 was used to find differentially abundant taxa among treatments.

## Acknowledgements

This research was supported by a Faculty Seed Grant from the Office of Research and Economic Development at the University of Nebraska-Lincoln. Dr. Andrew K. Benson, Mallory Van Haute, and Qinnan Yang were instrumental for sequencing library preparation and machine use.

## Author Contributions

ROC and CEC designed the study. ROC and JMV performed the experiments. ROC and CEC wrote the manuscript, and all authors approved the manuscript in its final form.

