## Supplemental Figures and Tables for "Targeted manipulation of abundant and rare taxa in the *Daphnia magna* microbiota with antibiotics impacts host fitness differentially"

| Microbiome PERMANOVA pairwise comparisons against NONE | | | | |  |  |
| --- | --- | --- | --- | --- | --- | --- |
| AZT | Df | SumsOfSqs | MeanSqs | F.Model | R2 | Pr (>F) |
| Antibiotic | 1 | 0.37472 | 0.37472 | 1.8969 | 0.13649 | 0.126 |
| Residuals | 12 | 2.37065 | 0.19755 |  | 0.86351 |  |
| ERY |  |  |  |  |  |  |
| **Antibiotic** | 1 | 0.7732 | 0.77322 | 3.6411 | 0.1764 | **0.01** |
| Residuals | 17 | 3.6101 | 0.21236 |  | 0.8236 |  |
| SFX |  |  |  |  |  |  |
| **Antibiotic** | 1 | 1.1052 | 1.10518 | 5.4953 | 0.28188 | **0.001** |
| Residuals | 14 | 2.8156 | 0.20111 |  | 0.71812 |  |
| ALL |  |  |  |  |  |  |
| **Antibiotic** | 1 | 1.228 | 1.228 | 6.2536 | 0.30876 | **0.001** |
| Residuals | 14 | 2.7492 | 0.19637 |  | 0.69124 |  |

Supplementary Table 1: Pairwise comparisons of the *Daphnia magna* microbiome across antibiotic treatments as compared to the no antibiotic control.

| Bacterial genera differing across antibiotic treatments | | |  |  |  |  |  |  |
| --- | --- | --- | --- | --- | --- | --- | --- | --- |
| AZT | Class | Genus | Base Mean | Log2 Fold Change | lfcSE | stat | p value | p adj |
|  | Flavobacteriia | Flavobacterium | 1298.24 | 4.66 | 1.3 | 3.58 | 0.00034151 | **0.00648867** |
|  | Alphaproteobacteria | Ensifer | 394.86 | 2.55 | 0.78 | 3.27 | 0.00105875 | **0.01609304** |
|  | Gammaproteobacteria | Pseudomonas | 417.82 | -26.51 | 4.73 | -5.6 | 2.09E-08 | **7.94E-07** |
|  | - | SR1 genera incertae sedis | 74.81 | -11.28 | 2.14 | -5.27 | 1.35E-07 | **3.41E-06** |
|  | Alphaproteobacteria | Sphingomonas | 23.24 | -24.29 | 2.73 | -8.87 | 7.17E-19 | **5.45E-17** |
|  | Alphaproteobacteria | Sphingomonas | 63.83 | 5.73 | 1.83 | 3.12 | 0.00177886 | **0.01868399** |
|  | Alphaproteobacteria | Microvirga | 15.78 | 7.04 | 2.28 | 3.1 | 0.00196674 | **0.01868399** |
|  | Gammaproteobacteria | Perlucidibaca | 14.83 | -8.13 | 2.57 | -3.16 | 0.00156232 | **0.01868399** |
| ERY | Class | Genus | Base Mean | Log2 Fold Change | lfcSE | stat | p value | p adj |
|  | Betaproteobacteria | Vitreoscilla | 1514.77 | -5.07 | 1.39 | -3.64 | 0.00027304 | **0.00207511** |
|  | Alphaproteobacteria | Sphingopyxis | 1350.47 | 2.90 | 0.88 | 3.29 | 0.00100825 | **0.00589437** |
|  | Betaproteobacteria | - | 777.00 | 3.21 | 0.99 | 3.26 | 0.0011253 | **0.00610878** |
|  | Sphingobacteriia | Pedobacter | 1008.73 | -8.48 | 2.12 | -4.00 | 6.42E-05 | **0.00069712** |
|  | Alphaproteobacteria | - | 513.49 | 1.87 | 0.69 | 2.71 | 0.00675451 | **0.03208392** |
|  | Cytophagia | Emticicia | 362.58 | -3.96 | 1.04 | -3.80 | 0.00014408 | **0.00121667** |
|  | Alphaproteobacteria | Ensifer | 394.86 | 2.71 | 0.78 | 3.48 | 0.00049934 | **0.00341211** |
|  | Sphingobacteriia | - | 186.05 | 20.18 | 4.76 | 4.24 | 2.23E-05 | **0.00042416** |
|  | Verrucomicrobiae | Prosthecobacter | 493.32 | -6.17 | 1.14 | -5.41 | 6.22E-08 | **2.36E-06** |
|  | Alphaproteobacteria | Sphingopyxis | 320.44 | 6.25 | 1.24 | 5.06 | 4.24E-07 | **1.08E-05** |
|  | - | SR1 genera incertae sedis | 74.82 | -7.58 | 1.98 | -3.82 | 0.00013321 | **0.00121667** |
|  | Flavobacteriia | Fluviicola | 304.89 | -7.86 | 1.90 | -4.13 | 3.62E-05 | **0.00055057** |
|  | Cytophagia | Emticicia | 96.78 | -9.43 | 2.73 | -3.46 | 0.00053875 | **0.00341211** |
|  | Alphaproteobacteria | - | 282.16 | -7.07 | 1.75 | -4.05 | 5.18E-05 | **0.00065577** |
|  | Alphaproteobacteria | Sphingomonas | 63.83 | 9.89 | 1.83 | 5.42 | 5.96E-08 | **2.36E-06** |
|  | Alphaproteobacteria | Microvirga | 15.78 | 6.29 | 2.28 | 2.76 | 0.00574213 | **0.02909346** |
| SFX | Class | Genus | Base Mean | Log2 Fold Change | lfcSE | stat | p value | p adj |
|  | Betaproteobacteria | Vitreoscilla | 1514.77 | -7.23 | 1.40 | -5.18 | 2.27E-07 | **5.21E-06** |
|  | Flavobacteriia | Flavobacterium | 1298.24 | 4.15 | 1.30 | 3.19 | 0.00142255 | **0.01635929** |
|  | Sphingobacteriia | Pedobacter | 1008.73 | -9.19 | 2.13 | -4.32 | 1.54E-05 | **0.00026632** |
|  | Sphingobacteriia | - | 186.05 | 27.43 | 4.75 | 5.77 | 7.84E-09 | **5.41E-07** |
|  | Alphaproteobacteria | Sphingopyxis | 320.44 | 6.71 | 1.24 | 5.43 | 5.59E-08 | **1.93E-06** |
|  | Betaproteobacteria | - | 204.07 | 2.06 | 0.73 | 2.85 | 0.00440856 | **0.04276069** |
|  | Alphaproteobacteria | Sphingomonas | 63.83 | 7.52 | 1.83 | 4.12 | 3.86E-05 | **0.00053231** |
|  | Alphaproteobacteria | Microvirga | 15.78 | 6.40 | 2.28 | 2.81 | 0.00495776 | **0.04276069** |
| ALL | Class | Genus | Base Mean | Log2 Fold Change | lfcSE | stat | p value | p adj |
|  | Betaproteobacteria | Vitreoscilla | 1514.77 | -4.18 | 1.47 | -2.85 | 0.00440375 | **0.02231235** |
|  | Alphaproteobacteria | Sphingopyxis | 1350.47 | 2.80 | 0.93 | 3.01 | 0.0026375 | **0.01670416** |
|  | Betaproteobacteria | - | 777.00 | 3.67 | 1.04 | 3.54 | 0.00040617 | **0.00340772** |
|  | Sphingobacteriia | Pedobacter | 1008.73 | -8.74 | 2.25 | -3.88 | 0.00010299 | **0.0013046** |
|  | Alphaproteobacteria | - | 513.49 | 2.41 | 0.73 | 3.32 | 0.00091197 | **0.00630086** |
|  | Cytophagia | Emticicia | 362.58 | -3.01 | 1.10 | -2.73 | 0.00629079 | **0.02988124** |
|  | Betaproteobacteria | Hydrogenophaga | 492.79 | 3.50 | 0.74 | 4.75 | 1.99E-06 | **3.79E-05** |
|  | Sphingobacteriia | - | 186.05 | 19.24 | 5.03 | 3.83 | 0.0001301 | **0.00141249** |
|  | Verrucomicrobiae | Prosthecobacter | 493.32 | -7.62 | 1.27 | -5.99 | 2.09E-09 | **1.59E-07** |
|  | Alphaproteobacteria | Sphingopyxis | 320.44 | 6.85 | 1.30 | 5.28 | 1.32E-07 | **4.51E-06** |
|  | - | SR1 genera incertae sedis | 74.82 | -6.08 | 2.08 | -2.93 | 0.00342309 | **0.01947719** |
|  | Betaproteobacteria | - | 204.07 | 1.96 | 0.76 | 2.57 | 0.01018991 | **0.04302407** |
|  | Flavobacteriia | Fluviicola | 304.89 | -7.58 | 2.02 | -3.75 | 0.00017654 | **0.00167714** |
|  | Alphaproteobacteria | - | 282.16 | -5.34 | 1.83 | -2.91 | 0.0035879 | **0.01947719** |
|  | Flavobacteriia | Flavobacterium | 146.45 | -5.54 | 2.21 | -2.51 | 0.01196684 | **0.04786738** |
|  | Alphaproteobacteria | Sphingomonas | 63.83 | 7.65 | 1.90 | 4.03 | 5.60E-05 | **0.00085058** |
|  | Flavobacteriia | Flavobacterium | 50.82 | -8.60 | 2.45 | -3.51 | 0.00044838 | **0.00340772** |
|  | Gammaproteobacteria | Perlucidibaca | 14.83 | -7.31 | 2.73 | -2.68 | 0.00740424 | **0.03310132** |
|  | Alphaproteobacteria | Caulobacter | 17.53 | -20.18 | 3.86 | -5.22 | 1.78E-07 | **4.51E-06** |

Supplementary Table 2: Differentially abundant taxa in the *Daphnia magna* microbiome across antibiotic treatments as compared to the no antibiotic control.

| Cumulative reproduction ANOVA post-hoc Tukey HSD | | | |  |
| --- | --- | --- | --- | --- |
| Antibiotics | diff | lwr | upr | p adj |
| **AZT-NONE** | -7.0416667 | -11.033012 | -3.0503211 | **1.81E-05** |
| ERY-NONE | -0.6770833 | -4.6684289 | 3.31426222 | 0.99042008 |
| **SFX-NONE** | -7.4375 | -11.428846 | -3.4461545 | **4.82E-06** |
| **ALL-NONE** | -10.427083 | -14.418429 | -6.4357378 | **5.50E-11** |
| **ERY-AZT** | 6.36458333 | 2.37323778 | 10.3559289 | **0.00015077** |
| SFX-AZT | -0.3958333 | -4.3871789 | 3.59551222 | 0.99880448 |
| ALL-AZT | -3.3854167 | -7.3767622 | 0.60592888 | 0.13949652 |
| **SFX-ERY** | -6.7604167 | -10.751762 | -2.7690711 | **4.47E-05** |
| **ALL-ERY** | -9.75 | -13.741346 | -5.7586545 | **6.61E-10** |
| ALL-SFX | -2.9895833 | -6.9809289 | 1.00176222 | 0.2434544 |
| Temperature |  |  |  |  |
| **11C-19C** | -38.575 | -40.386427 | -36.763573 | **2.24E-11** |
| Antibiotics:Temperature |  |  |  |  |
| **AZT:19C-NONE:19C** | -14.354167 | -20.907015 | -7.8013182 | **5.27E-10** |
| ERY:19C-NONE:19C | -1.3541667 | -7.9070151 | 5.19868175 | 0.99970775 |
| **SFX:19C-NONE:19C** | -14.875 | -21.427848 | -8.3221516 | **1.19E-10** |
| **ALL:19C-NONE:19C** | -20.854167 | -27.407015 | -14.301318 | **2.25E-11** |
| **NONE:11C-NONE:19C** | -48.916667 | -55.469515 | -42.363818 | **2.24E-11** |
| **AZT:11C-NONE:19C** | -48.645833 | -55.198682 | -42.092985 | **2.24E-11** |
| **ERY:11C-NONE:19C** | -48.916667 | -55.469515 | -42.363818 | **2.24E-11** |
| **SFX:11C-NONE:19C** | -48.916667 | -55.469515 | -42.363818 | **2.24E-11** |
| **ALL:11C-NONE:19C** | -48.916667 | -55.469515 | -42.363818 | **2.24E-11** |
| **ERY:19C-AZT:19C** | 13 | 6.44715158 | 19.5528484 | **2.95E-08** |
| SFX:19C-AZT:19C | -0.5208333 | -7.0736818 | 6.03201509 | 0.99999993 |
| ALL:19C-AZT:19C | -6.5 | -13.052848 | 0.05284842 | 0.05391977 |
| **NONE:11C-AZT:19C** | -34.5625 | -41.115348 | -28.009652 | **2.24E-11** |
| **AZT:11C-AZT:19C** | -34.291667 | -40.844515 | -27.738818 | **2.24E-11** |
| **ERY:11C-AZT:19C** | -34.5625 | -41.115348 | -28.009652 | **2.24E-11** |
| **SFX:11C-AZT:19C** | -34.5625 | -41.115348 | -28.009652 | **2.24E-11** |
| **ALL:11C-AZT:19C** | -34.5625 | -41.115348 | -28.009652 | **2.24E-11** |
| **SFX:19C-ERY:19C** | -13.520833 | -20.073682 | -6.9679849 | **6.43E-09** |
| **ALL:19C-ERY:19C** | -19.5 | -26.052848 | -12.947152 | **2.25E-11** |
| **NONE:11C-ERY:19C** | -47.5625 | -54.115348 | -41.009652 | **2.24E-11** |
| **AZT:11C-ERY:19C** | -47.291667 | -53.844515 | -40.738818 | **2.24E-11** |
| **ERY:11C-ERY:19C** | -47.5625 | -54.115348 | -41.009652 | **2.24E-11** |
| **SFX:11C-ERY:19C** | -47.5625 | -54.115348 | -41.009652 | **2.24E-11** |
| **ALL:11C-ERY:19C** | -47.5625 | -54.115348 | -41.009652 | **2.24E-11** |
| ALL:19C-SFX:19C | -5.9791667 | -12.532015 | 0.57368175 | 0.10823123 |
| **NONE:11C-SFX:19C** | -34.041667 | -40.594515 | -27.488818 | **2.24E-11** |
| **AZT:11C-SFX:19C** | -33.770833 | -40.323682 | -27.217985 | **2.24E-11** |
| **ERY:11C-SFX:19C** | -34.041667 | -40.594515 | -27.488818 | **2.24E-11** |
| **SFX:11C-SFX:19C** | -34.041667 | -40.594515 | -27.488818 | **2.24E-11** |
| **ALL:11C-SFX:19C** | -34.041667 | -40.594515 | -27.488818 | **2.24E-11** |
| **NONE:11C-ALL:19C** | -28.0625 | -34.615348 | -21.509652 | **2.24E-11** |
| **AZT:11C-ALL:19C** | -27.791667 | -34.344515 | -21.238818 | **2.24E-11** |
| **ERY:11C-ALL:19C** | -28.0625 | -34.615348 | -21.509652 | **2.24E-11** |
| **SFX:11C-ALL:19C** | -28.0625 | -34.615348 | -21.509652 | **2.24E-11** |
| **ALL:11C-ALL:19C** | -28.0625 | -34.615348 | -21.509652 | **2.24E-11** |
| AZT:11C-NONE:11C | 0.27083333 | -6.2820151 | 6.82368175 | 1 |
| ERY:11C-NONE:11C | 8.70E-14 | -6.5528484 | 6.55284842 | 1 |
| SFX:11C-NONE:11C | 9.59E-14 | -6.5528484 | 6.55284842 | 1 |
| ALL:11C-NONE:11C | -4.00E-14 | -6.5528484 | 6.55284842 | 1 |
| ERY:11C-AZT:11C | -0.2708333 | -6.8236818 | 6.28201509 | 1 |
| SFX:11C-AZT:11C | -0.2708333 | -6.8236818 | 6.28201509 | 1 |
| ALL:11C-AZT:11C | -0.2708333 | -6.8236818 | 6.28201509 | 1 |
| SFX:11C-ERY:11C | 8.88E-15 | -6.5528484 | 6.55284842 | 1 |
| ALL:11C-ERY:11C | -1.27E-13 | -6.5528484 | 6.55284842 | 1 |
| ALL:11C-SFX:11C | -1.36E-13 | -6.5528484 | 6.55284842 | 1 |

Supplementary Table 3: Tukey HSD comparisons of cumulative *Daphnia magna* reproduction across antibiotic treatments.

| Day of first brood ANOVA post-hoc Tukey HSD | | | |  |
| --- | --- | --- | --- | --- |
| *Antibiotics* | diff | lwr | upr | p adj |
| AZT-ALL | 0.25430598 | -0.6235563 | 1.1321683 | 0.9313473 |
| ERY-ALL | 0.21130952 | -0.6622268 | 1.0848458 | 0.963571 |
| NONE-ALL | -0.431746 | -1.3187666 | 0.4552745 | 0.6673479 |
| **SFX-ALL** | 0.92380952 | 0.03678896 | 1.8108301 | **0.0366059** |
| ERY-AZT | -0.0429965 | -0.8913897 | 0.8053968 | 0.9999145 |
| NONE-AZT | -0.686052 | -1.5483228 | 0.1762188 | 0.1879584 |
| SFX-AZT | 0.66950355 | -0.1927673 | 1.5317744 | 0.2088342 |
| NONE-ERY | -0.6430556 | -1.5009218 | 0.2148107 | 0.2405979 |
| SFX-ERY | 0.7125 | -0.1453662 | 1.5703662 | 0.1538504 |
| **SFX-NONE** | 1.35555556 | 0.48396262 | 2.2271485 | **0.00027** |

Supplementary Table 4: Tukey HSD comparisons of *Daphnia magna* day of first brood across antibiotic treatments.

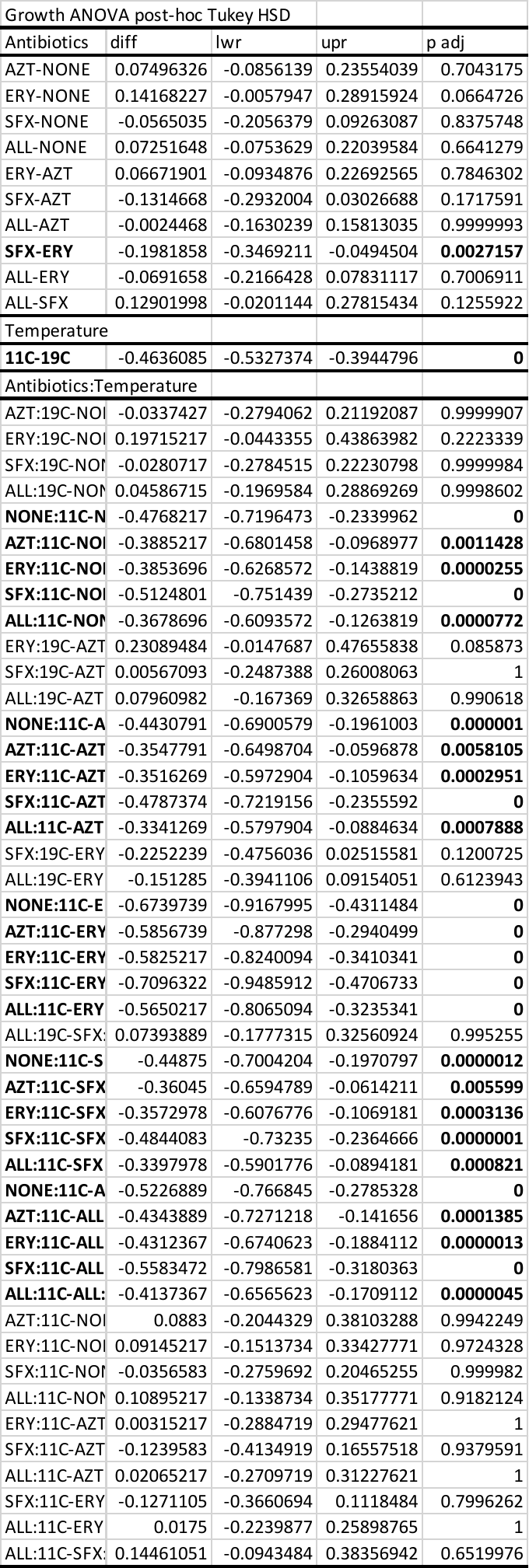

Supplementary Table 5: Tukey HSD comparisons of *Daphnia magna* growth across treatments.

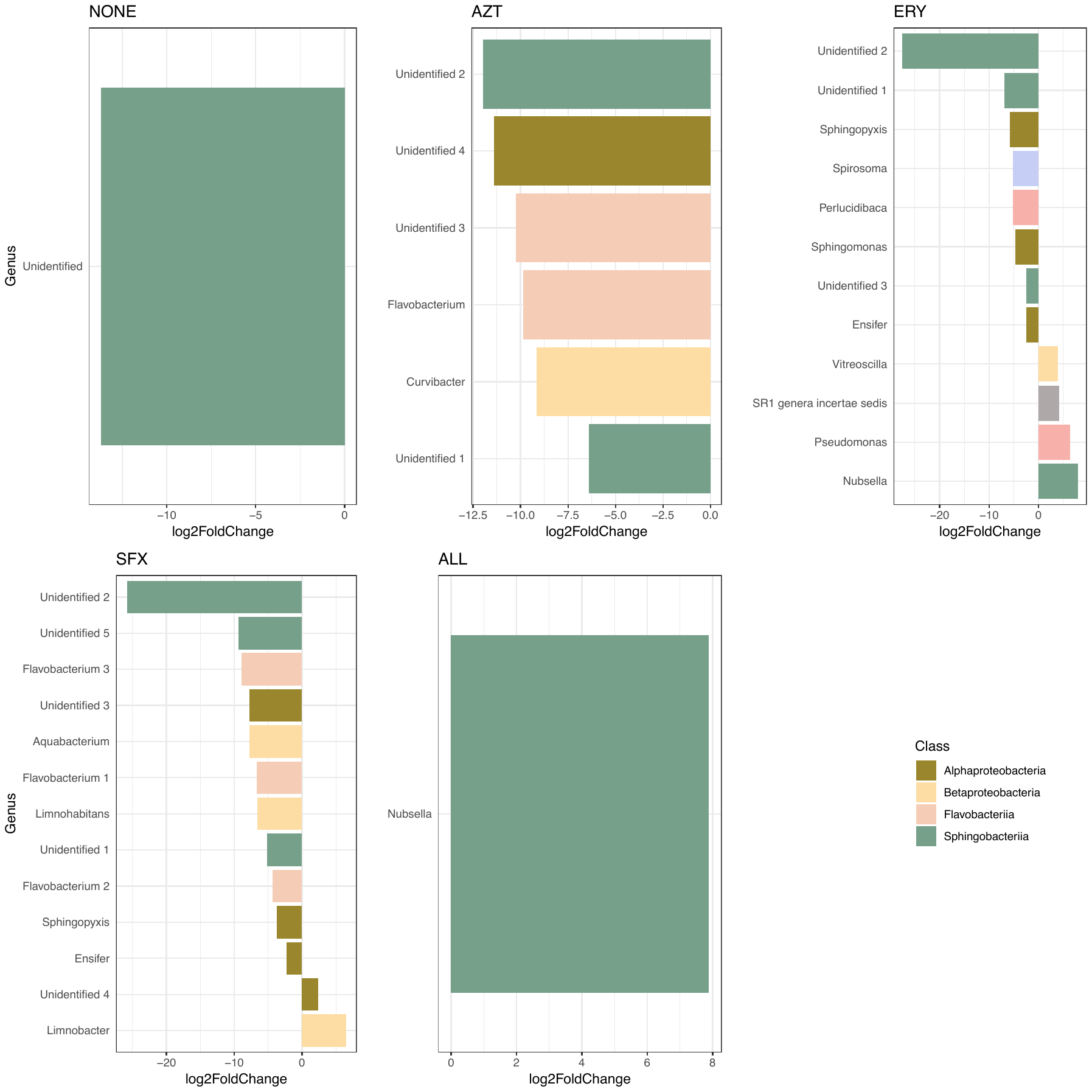

Supplementary Figure 1: Differentially abundant ASVs in each antibiotic treatment in 11°C as compared to the same antibiotic treatment in 19°C. Each bar represents a single ASV identified to the genus level, with genus name indicated on the left. Bar color indicates the bacterial class of each ASV, and bar length indicates the fold change in abundance of each ASV.

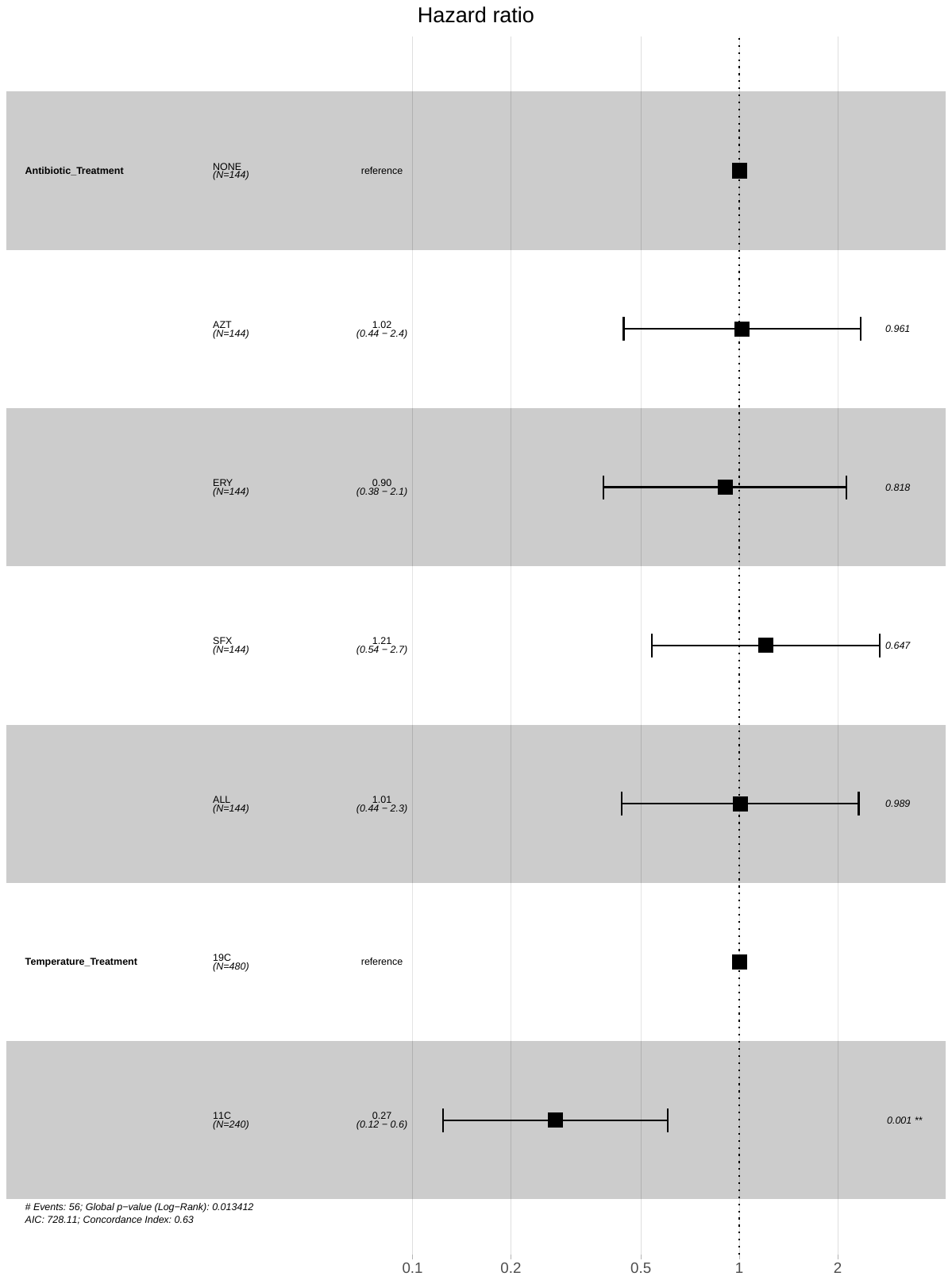

Supplementary Figure 2: *Daphnia magna* survival hazard ratios across antibiotic treatments and temperatures.
